# Ketamine disrupts consciousness in healthy participants in relation with psychotic-like symptoms

**DOI:** 10.1101/2025.11.06.687042

**Authors:** Lucie Berkovitch, Alexandre Salvador, Thomas Andrillon, Stanislas Dehaene, Raphaël Gaillard

## Abstract

Ketamine is an NMDA-receptor antagonist, which alters the state of wakeful consciousness at high doses. At lower doses, it induces reversible psychotic-like symptoms and has been used as a pharmacological model of psychosis. In this study, we explore whether low doses of ketamine disrupt conscious access in relation with its psychotomimetic effects, and investigate the neural correlates of its action.

We administered ketamine and placebo to 21 healthy volunteers following a double-blind within-subject randomized design and recorded brain activity with high-density electroencephalography during a perceptual task. Participants had to categorize a sound and a masked digit, and to report the digit visibility. To manipulate visibility, the delays between the sound, the digit, and the mask were varied. Principal component analysis was used to decompose ketamine-induced psychiatric symptoms and to examine their relationships with conscious access measures.

Under ketamine, participants had an increased visual masking effect, more interference between the sound and the digit, and a reduced ability to consciously perceive the digit. The N1 component, a EEG marker of visual processing, correlated with conscious access and was significantly reduced under ketamine. Ketamine induced manic-like and psychotic-like symptoms but only the psychotic-like dimension correlated with conscious access impairments.

Overall, our results suggest that ketamine disrupts conscious access in healthy subjects through an attenuation of early visual responses, and in relation with its psychotomimetic effects. Interestingly, these changes differed in part from those observed in patients with schizophrenia, opening new perspectives on the mechanisms of psychotic symptoms.

## Introduction

Across a variety of paradigms, a disruption of consciousness has been observed in persons with schizophrenia^1,2^. This disruption could play a role in the advent of psychotic episodes^1,3^ and correlates with the intensity of positive symptoms^4^. Ketamine is a noncompetitive *N*-methyl-D-aspartate receptor antagonist that is used in medicine as an anesthetic^5^, inducing a rapid loss of consciousness when administered at high doses^2,36,7^. At lower doses, it can trigger reversible psychotic-like symptoms such as delusional ideas in healthy subjects and symptoms that mimic a relapse in patients with a remitted schizophrenia^8–11^. Given its behavioral, imaging and electrophysiological effects, it has been used as a pharmacological model of early psychosis^12–15^. However, whether psychotomimetic doses of ketamine disrupt consciousness remains unknown. More broadly, the relationships between a disruption of consciousness and psychotic symptoms have never been causally studied.

In this context, the goal of the present project was to evaluate whether low doses of ketamine impair conscious access and, if so, by which mechanisms. In particular, we aim to distinguishing between sensory evidence accumulation deficits and attentional alterations. Indeed, according to the global neuronal workspace (GNW) theory^16,17^, conscious access rests upon the transient stabilization of neuronal activity encoding a specific piece of information amplified by top-down attention^18–20^. NMDA receptors have been shown to be involved in attentional amplification^21–23^. By blocking NMDA and upregulating AMPA receptors, ketamine could cause an increased feed-forward/feed-back imbalance^24^. Consistently, empirical data found that ketamine disrupts a range of predictive top-down cognitive and perceptual processing, such as prior-informed visual recognition^25^, figure-ground modulation^21,26^ and balance between external and internal modes^27^. Other studies suggest that ketamine disrupts both feed-forward and feed-back processing^28^. Electrophysiologically, ketamine reduces several ERP components, including N2, P2, P3, processing negativity and mismatch negativity, whereas other components remain generally unaffected (P50, PPI, P1, and N1)^29^. Thus, ketamine may alter consciousness through its effects on event-related potentials in the 200-500 ms time window (N2, P3) which have been related to a fast non-linear ignition at the onset of conscious perception^30–35^.

In the current study, we used a variant of previous paradigms exploring conscious access and its disruption in patients with schizophrenia^2,3,36,37^. We administered an intravenous continuous infusion of ketamine or placebo to healthy volunteers in a cross-over, randomized, double-blind paradigm and recorded brain activity using high-resolution electroencephalography. Participants performed a perceptual task manipulating two ways of disrupting conscious access: metacontrast masking (which impairs visual perception through competition between two visual stimuli presented in brief succession) and attentional distraction (which corners attentional central resources, here by an auditory stimulus).

We first hypothesized that ketamine would impair conscious access. Second, we expected to observe a relationship between consciousness disruption and ketamine-induced psychotomimetic effects. At the neural level, we explored whether ketamine would affect early visual evoked potentials, late responses, or perhaps both of them.

## Results

### Ketamine-induced symptoms and ketamine blood levels

Healthy adult participants (n = 21) were tested twice at least one month apart, following a double-blind, placebo-controlled, randomized crossover design. They received either ketamine (maximum 1 mg/kg over 2h) or placebo (sodium chloride) intravenously. Psychiatric symptoms measurements were conducted 30 minutes after the infusion started with the Clinician-Administered Dissociative States Scale (CADSS) capturing dissociation, and Brief Psychiatric Rating Scale (BPRS) measuring psychiatric-like symptoms. Ketamine blood levels were controlled right before clinical assessment.

On ketamine visits, CADSS scores ranged from 0 to 45 (mean: 15.52, SD: 11.66) and BPRS scores ranged from 24 – which is the minimal score – to 41 (mean: 30.62, SD: 5.28), see Figure 1A, left.

**Figure 1.**
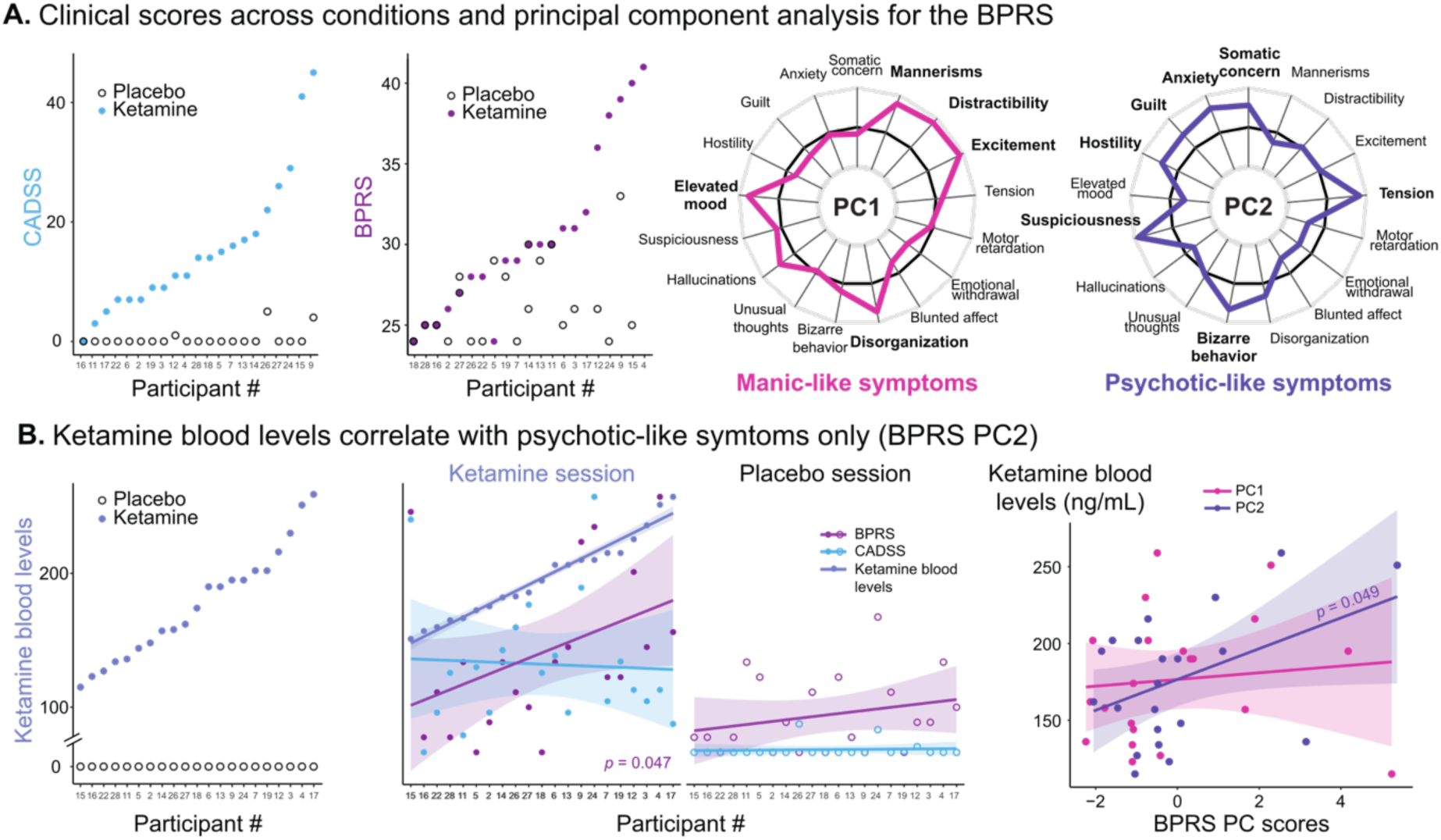
Ketamine-induced symptoms and ketamine blood levels. A. Clinical scores across visits and principal component analysis for the BPRS. Left. CADSS (light blue) and BPRS (deep magenta) for each participant in each pharmacological condition. Each dot represents a subject in either the ketamine (colored) or the placebo (black outline) condition. Participants are ranked increasingly for each corresponding variable on the ketamine visits. There were important variations in clinical scores across subjects. Right. Results of the principal component analysis (PCA) for the BPRS items on the ketamine visits. Left. PCA yielded two significant principal components (PC) accounting for the 43% of the symptoms. Radar plots depict loadings of PC1 (pink) which represents manic-like symptoms and PC2 (purple) which represents psychotic-like symptoms. B. Ketamine blood levels correlate with BPRS PC2. Left. Ketamine blood levels (soft indigo) for each participant in each pharmacological condition. Each dot represents a subject in either the ketamine (colored) or the placebo (black outline) condition. Participants are ranked increasingly according to ketamine blood levels. There were important variations in ketamine blood levels across subjects. Middle. Ketamine blood levels (soft indigo), BPRS (deep magenta) and CADSS (light blue) were scaled between 0 and 1. Each dot represents a subject in either the ketamine (filled with color) or the placebo (outlined) condition, lines represent linear regressions, and shaded areas represent 95% confidence intervals. Participants are ranked increasingly as a function of ketamine blood level on the ketamine visits. Only BPRS scores significantly correlated with the ketamine blood levels. Right. Correlation between BPRS PC scores and ketamine blood levels. Each dot represents PC scores for each subject (PC1 pink, PC2 purple) as a function of ketamine blood levels, lines represent linear regressions, and shaded areas represent 95% confidence intervals. Only BPRS PC2 (purple) significantly correlated with ketamine blood levels.

To explore the dimensions of ketamine-induced psychiatric-like symptoms, we conducted a principal component analysis of the BPRS items. It yielded two significant principal components (PC) accounting for the 43% of symptoms variance. The first PC (BPRS PC1) explained 23.9% of the variance and loaded on manic-like items: excitement, distractibility, mannerisms, elevated mood, disorganization, and hallucinations. BPRS PC2 explained 18.9% of the variance and loaded on positive psychotic-like symptoms: suspiciousness, tension, bizarre behavior, anxiety, somatic concern, guilt, hostility, disorganization (Figure 1A, right).

On ketamine visits, ketamine blood levels ranged from 115 ng/mL to 259 ng/mL (mean: 1740, SD: 41.52, see Figure 1B, left). Ketamine blood levels significantly correlated with BPRS total scores (Pearson *r* = 0.44, *t_19_* = 2.12, *p* = 0.047) and BPRS PC2 (*r* = 0.43, *t_19_* = 2.10, *p* = 0.0497) but not with other clinical measures (BPRS PC1: *r* = 0.11, *t_19_* = 0.46, *p* = 0.65, 1/BF = 2.0; CADSS: *r* = −0.10, *t_19_* = −0.45, *p* = 0.66, 1/BF = 1.99), see Figure 1B.

### Behavioral measures: ketamine disrupts conscious access

Participants performed a visual perceptual task designed to disentangle attentional and sensory contributions to conscious access. At the start of each trial, a sound was played. After a variable delay, a digit target (or a catch) was briefly presented, followed, after another variable delay, by a metacontrast mask. The main task was to indicate if the masked digit was greater or smaller than 5 and to report its visibility; the distracting task was to categorize the sound. Across different blocks, instructions required focusing either on the digit (single-task), or on the sound (unattended condition), or on both (dual-task). Overall, our experimental design manipulated three variables: (1) stimulus relevance, by instructing participants to attend to the sound, the digit or both, (2) the amount of allocated attention in the dual task, by varying the sound-digit delay, (3) the amount of visual masking, by varying the digit-mask delay. The experimental paradigm is summarized in Figure 2 and detailed in the Methods section.

**Figure 2.**
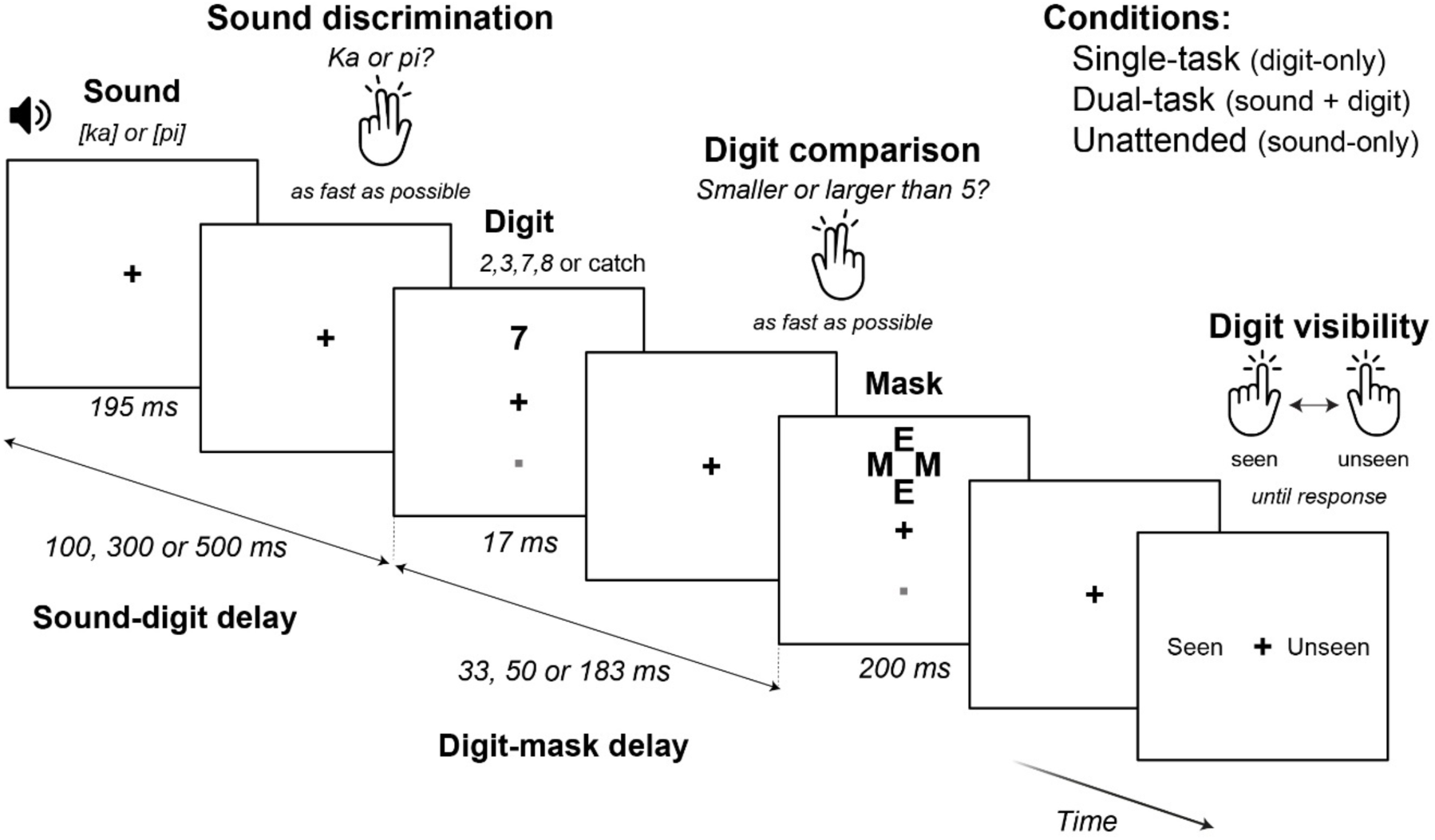
Experimental paradigm. A sound (an isolated syllable “ka” or “pi”) was played. After a first delay (sound-digit delay), a digit (or a catch in 20% of the trials) was briefly displayed (17 ms) above or below the fixation cross and subsequently masked after a second delay (digit-mask delay). The exact same sequences of stimuli were presented under three distinct conditions, which differed only in the requested task. In the dual-task condition, subjects were asked to: (1) determine as fast as possible whether the sound was “ka” or “pi”, (2) decide as fast as possible whether the digit was larger or smaller than 5 and (3) report the digit visibility using a categorical response “Seen” or “Unseen”. In the single-task condition, participants had to answer the two questions about the digit only (i.e., questions 2 and 3) and were asked to ignore the sound. Conversely, in the unattended condition, participants had to answer the question about the sound only (i.e., question 1) and to ignore the digit. Responses and reaction times were recorded.

To understand the respective effects of ketamine on masking effects and attentional availability, we conducted ANOVAs with different behavioral measures (discrimination, visibility and detection abilities measured with *d*-primes) as dependent variables, and task (single or dual task), sound-digit delay, digit-mask delay, and the pharmacological condition as within-subject variables. For clarity, we first describe the effect of the digit-mask delay (masking) and then of the sound-digit delay (attentional availability).

#### Increased masking effect

As expected, discrimination, visibility and detection abilities were significantly modulated by the digit-mask delay: the shorter the digit-mask delay, the lower discrimination, visibility and detection abilities (Figure 3). Importantly, the pharmacological condition also modulated these measures independently and in interaction with digit-mask delays.

**Figure 3.**
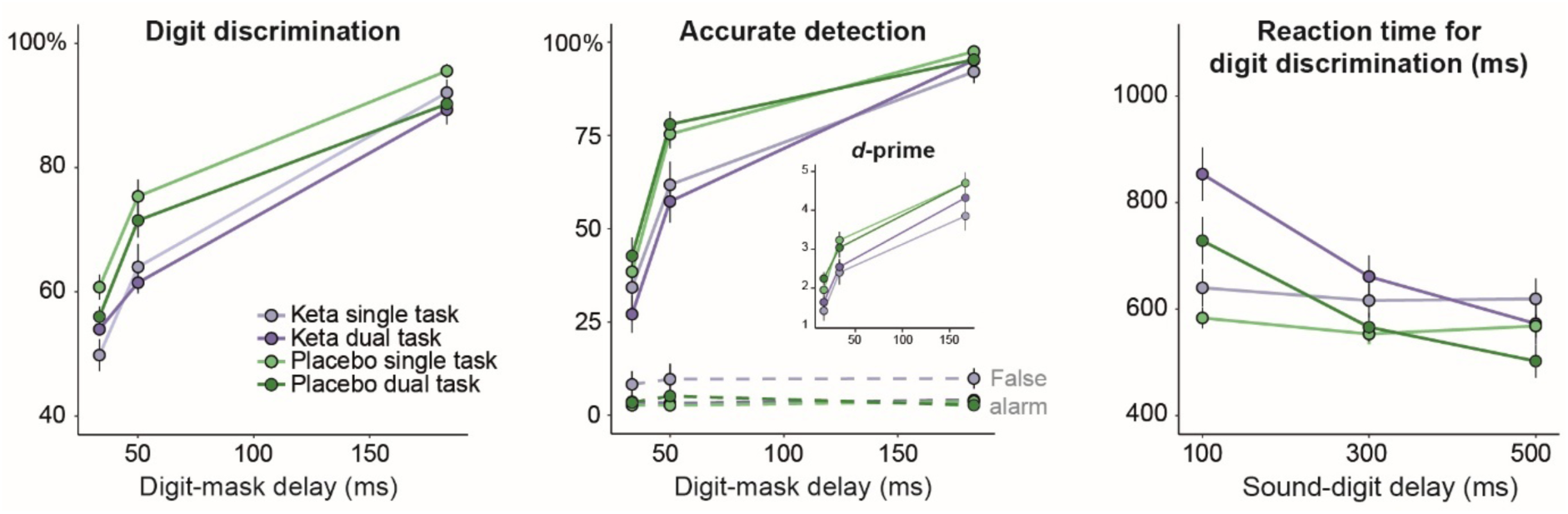
Ketamine increases masking sensitivity and interferes with attentional processes. Left. Performance in digit discrimination (i.e., comparing the masked digit to 5) as a function of digit-mask delays in the single-task and the dual-task conditions under placebo and ketamine (y scale = % correct). Error bars represent one standard error of the mean. Middle. Accurate detection (i.e., proportion of seen trials) for digit (plain lines) and catch trials (dashed lines) as a function of digit-mask delays in the single-task and the dual-task conditions under placebo and ketamine. The inset depicts detection *d*-primes, indexing how well visibility ratings distinguish between digit and catch trials. Error bars represent one standard error of the mean. Both for detection and discrimination, there was a main effect of digit-mask delays, task, and the pharmacological condition: ketamine increased sensitivity to masking. Right. Reaction times for digit discrimination as a function of digit-mask delays in the single-task and the dual-task conditions under placebo and ketamine. Error bars represent one standard error of the mean. In the dual-task condition, there was a significant interaction between the pharmacological condition and sound-digit delays, suggesting that ketamine interferes with attentional processes.

Specifically, for the discrimination task, i.e., the ability to compare the digit against 5, we found that performances increase with the digit-mask delay (*F_2,36_* = 233.40, *p* < 0.001), and decrease under ketamine (*F_2,18_* = 38.96, *p* < 0.001) with a significant interaction between the digit -digit delay and the pharmacological condition (*F_2,36_* = 5.40, *p* = 0.009; see Figure 3, left). There was a main effect of the task context (higher performances in single versus dual task; *F_1,18_* = 6.25, *p* = 0.022) which did not interact with the digit-mask delay (*F_2,36_* = 1.67, *p* = 0.20). When analyzing the two contexts separately, there was a main effect of the digit-mask delay and the pharmacological condition in both dual and single task contexts (all *p* < 0.01), but their interaction was significant in the dual task only (*F_2,36_* = 10.58, *p* < 0.001; single task: *F_2,36_* = 2.40, *p* = 0.11). Indeed, in the dual-task, ketamine-induced alterations predominate at the intermediate digit-mask delay (50 ms: *F_1,18_* = 24.61, *p* < 0.001; *p* > 0.1 at other digit-mask delays). A different pattern was observed for the single-task, where ketamine-induced alterations occurred both at the intermediate and the short digit-mask delays (33 ms: *F_1,18_* = 22.54, *p* < 0.001; 50 ms: *F_1,18_* = 8.90, *p* = 0.008; 183 ms: *F_1,18_* = 3.40, *p* = 0.082).

Detection *d*-primes were computed by examining how subjective visibility reports (“Seen” versus “Unseen”) varied with the presence or absence of a target (digit versus catch trials). We found that performances increase with digit-mask delay (*F_2,36_* = 106.7, *p* < 0.001) and decrease under ketamine (*F_1,18_* = 6.83, *p* = 0.018) with no significant interaction between the two, nor significant effect of task context (all *p* > 0.2, see Figure 3, middle). In both contexts, there was a main effect of digit-mask delay and pharmacological condition on *d*-primes (all *p* < 0.005), but no interaction between the two (all *p* > 0.7). Ketamine-induced alterations were observable at the short digit-mask delay in the dual-task (*F_1,18_* = 4.67, *p* = 0.045) and at the intermediate digit-mask delay in the single-task (*F_1,18_* = 4.87, *p* = 0.041).

Overall, we found increased masking effect under ketamine both for the single and the dual task.

#### Attentional interferences

To study ketamine effects on attention, we measured how sound-digit delays modulated reaction times (RT) in the digit discrimination task. Indeed, when two relevant stimuli have to be processed in close succession, there is a slowdown in the processing of the second stimulus (here, the digit) because attention is still focused on the first one (here, the sound)^38,39^. This phenomenon is called the psychological refractory period (PRP) and is tightly related to the attentional blink (where conscious perception of the second stimulus vanishes).

In our paradigm, we observed a PRP in the dual-task condition with a strong effect of the sound-digit delay on RT (*F_2,36_* = 92.84, *p* < 0.001; see Figure 4, right). We also observed a more pronounced slowdown under ketamine compared to placebo (*F_1,18_* = 7.71, *p* = 0.013), and a significant interaction between the sound-digit delay and pharmacological condition (*F_2,36_* = 5.28, *p* = 0.010, see Figure 3, right). Additional results regarding attentional interferences can be found in the Supplementary Materials.

**Figure 4.**
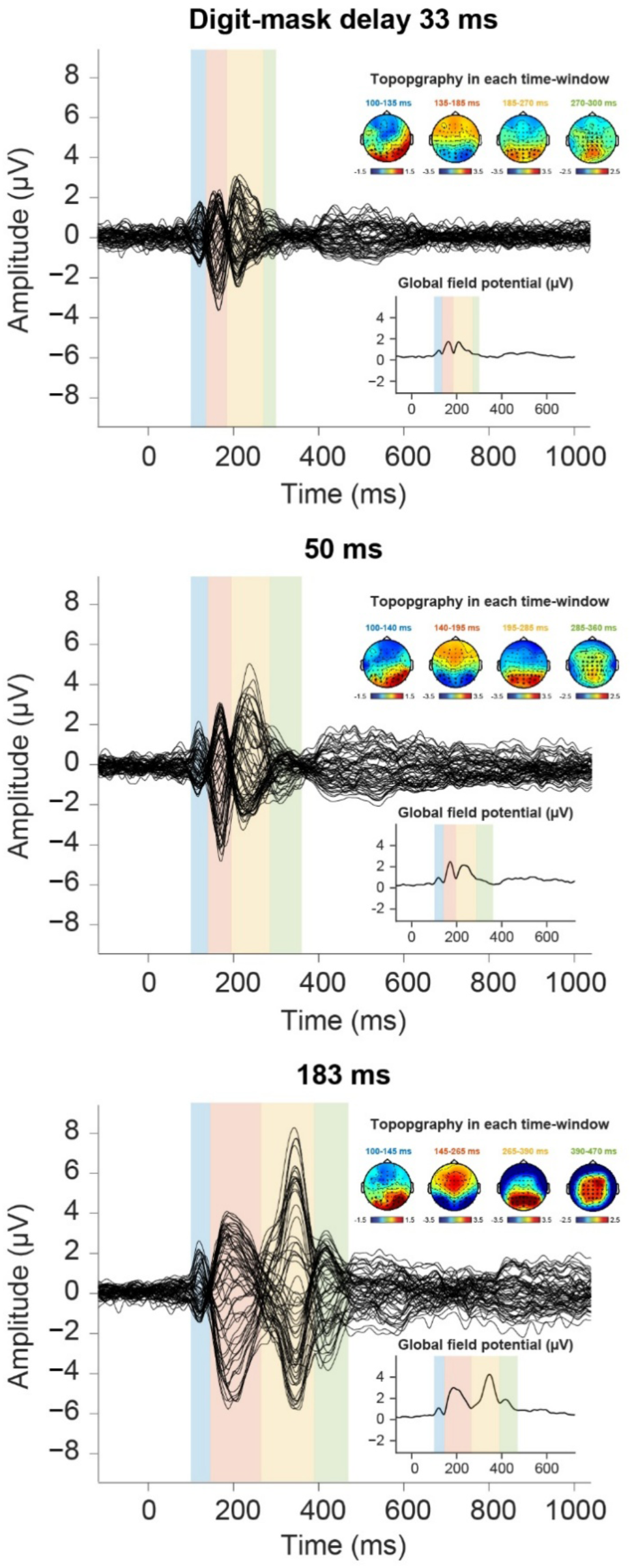
Time-course of brain activity. Each subplot shows the time course of mask-subtracted ERP for each digit-mask delay in the dual-task condition (averaged across participants and pharmacological conditions). Global field potentials (GFP) are shown in inserts. Specific time-windows were determined according to GFP for each digit-mask delays and are shown by colored rectangles. EEG voltage topographies are shown for each time-windows. Clusters of electrodes for each time-widows were determined to capture components at the longest digit-mask delay (183 ms). They are depicted by black dots in the topographies for each time-windows and were kept the same across digit-mask delays.

Overall, we found that ketamine modulated the PRP, suggesting that it interfered with attentional processes.

#### Poorer performances in the sound-related task

Participants were overall less able to perform the sound-related task under ketamine (see Supplementary Materials for details).

### EEG measures: ketamine reduces the amplitude of ERPs tracking conscious access

We then turn to event-related potentials (ERPs) evoked by the digit, in order to uncover the brain mechanisms through which ketamine impairs conscious access. Event-related potentials (ERP) components were identified based on latency, topographical responses and previous work^21,28^. ERP latency highly varied according to the digit-mask delays, so different time-windows were used for each digit-mask delay (see Figure 4 and Methods for details on potential definition)

#### N1 and P3 increase as a function of digit-mask delays but only P3 depends on task-relevance

As a reference, we first investigated the effects of masking and attention for each potential in the placebo condition. Across all tasks, several components increased as a function of digit-mask delays: N1 (*F_1,17_* = 5.82, *p* = 0.027), “early” P3 (*F_1,17_* = 12.79, *p* = 0.002), and “late” P3 (*F_1,17_* = 24.73, *p* < 0.001). Both early and late P3 components were modulated by the task context (early P3: task effect: *F_2,34_* = 7.09, *p* = 0.003, interaction task × digit-mask delay: *F_2,34_* = 4.52, *p* = 0.018; late P3: task effect: *F_2,34_* = 10.20, *p* < 0.001; interaction task × digit-mask delay: *p* = 0.8) whereas N1 was not (task effect and interaction task × digit-mask delay: all *p* > 0.4). Specifically, early and late P3 amplitudes were larger in the attended (single and dual tasks) compared to the unattended (sound-only) conditions (early P3: *F_1,17_* = 10.39, *p* = 0.005, late P3: *F_1,17_* = 24.73, *p* < 0.001). This suggests that P3 depends on task-relevant top-down processes whereas N1 did not.

In line with the absence of its effects on discrimination or visibility performances, sound-digit delays had only a minor impact on ERP components (see Supplementary Materials for details). Therefore, sound-target delays were collapsed for the subsequent analyses.

#### Ketamine alters N1

In the dual-task condition, ketamine significantly decreased the N1 component amplitude (*F_1,17_* = 5.84, *p* = 0.027), without significantly interacting with digit-mask delays (see Figure 5, left). There was no other significant effect of the pharmacological condition (all *p* > 0.3).

**Figure 5.**
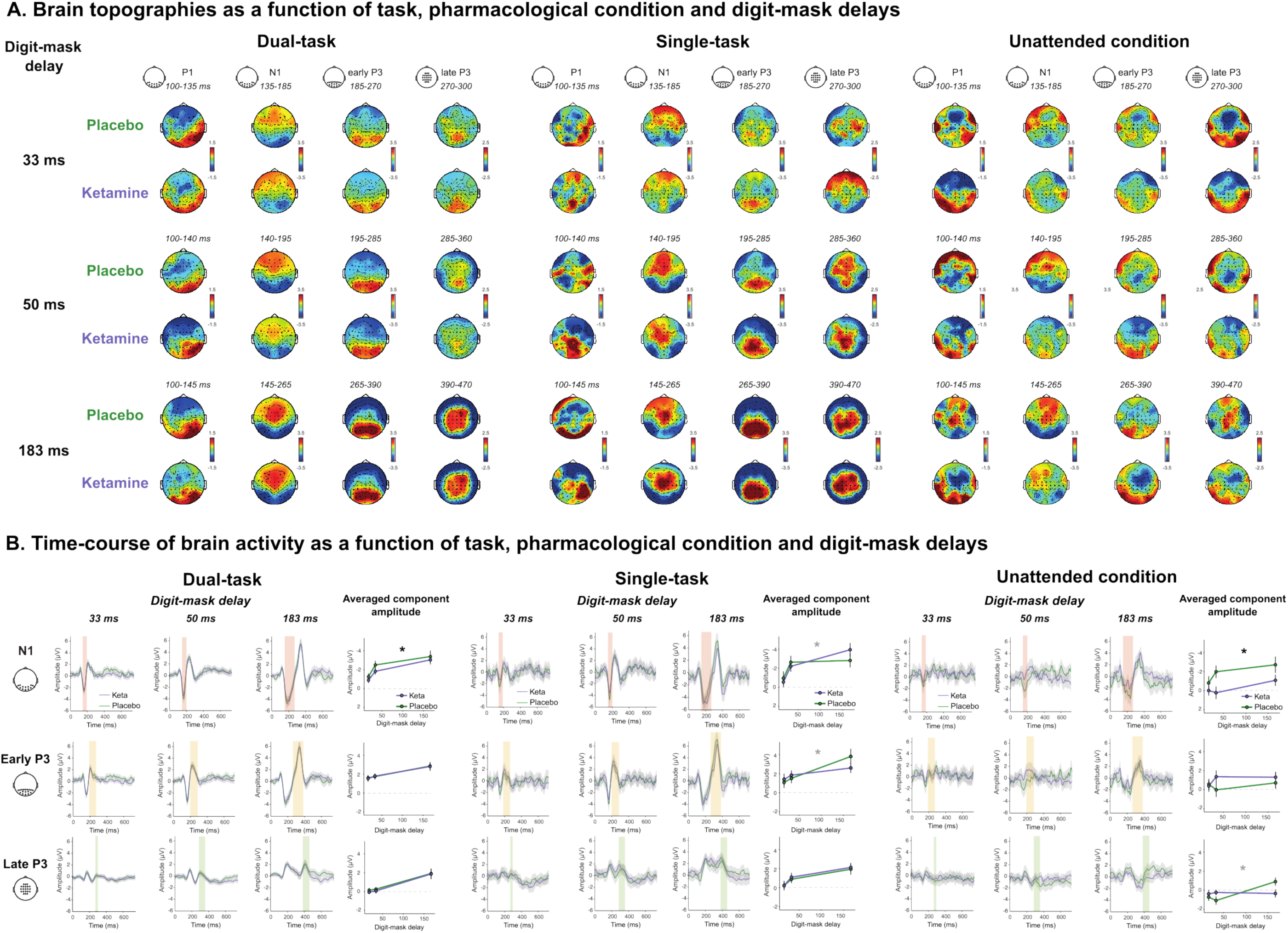
Ketamine alters brain activity as a function of tasks and digit-mask delays. A. Mask-subtracted ERP topographies for each potential time-window in the different task conditions (horizontally), i.e., dual-task vs. single-task vs. unattended condition, for each digit-mask delay in each pharmacological condition (vertically), i.e., placebo vs. ketamine. Small topographies on top depict the clusters of electrodes that were used to compute the averaged amplitude of each component. B. Time-course of brain-related activity under ketamine and placebo. Subplots show the time-course of ERPs in each condition of task, digit-mask delay and pharmacological condition (green: placebo, purple: ketamine). For each component, the preselected cluster of electrodes is depicted by black dots in the topographies at left. Preselected time-windows of interest, used for statistical analysis, are shown by colored rectangles. Grey shaded area around the curves represents one standard error of the mean. The line-plots depict the averaged amplitude of each component in this window, i.e., across the corresponding cluster of electrodes, as a function of digit-mask delays for each time-window in the different conditions. Error bars represent one standard error of the mean. * means p < 0.05 (black: significant difference between ketamine and placebo, gray: significant interaction between pharmacological condition and digit-mask delay).

In the single-task condition, there was no main effect of ketamine (all *p* > 0.3). However, ketamine interaction with digit-mask delays significantly modulated the N1 component amplitude (*F_1,17_* = 6.89, *p* = 0.018) and the early P3 component (*F_1,17_* = 5.46, *p* = 0.032, see Figure 5, middle).

In the unattended condition (sound-only), ketamine significantly increased P1 (*F_1,17_* = 5.82, *p* = 0.027) and decreased N1 amplitudes (*F_1,17_* = 11.35, *p* = 0.036). The interaction between the pharmacological condition and the digit-mask delays significantly modulated the late P3 amplitude (*F_1,17_* = 5.14, *p* = 0.037, see Figure 5, right).

Overall, N1 was decreased under ketamine both in the dual-task and the unattended condition, reflecting a modulation of early visual processing. Ketamine also modulated the P3 but in a less consistent fashion (no modulation in the dual-task but modulation in both single tasks)

#### N1 tracks conscious access

We then studied the relationship between EEG component amplitudes and behavioral consciousness measures. Since discrimination performances and visibility ratings were collected on a trial-by-trial basis, we separated conscious (i.e., seen and correctly discriminated) from unconscious (unseen) trials. Because most of the trials were conscious at the longest digit-mask delay (183 ms) and unconscious at the shortest digit-mask delay (33 ms), as shown in Figure 2, we restricted the following analyses to the intermediate (50 ms) digit-mask delay (Figure 6).

**Figure 6.**
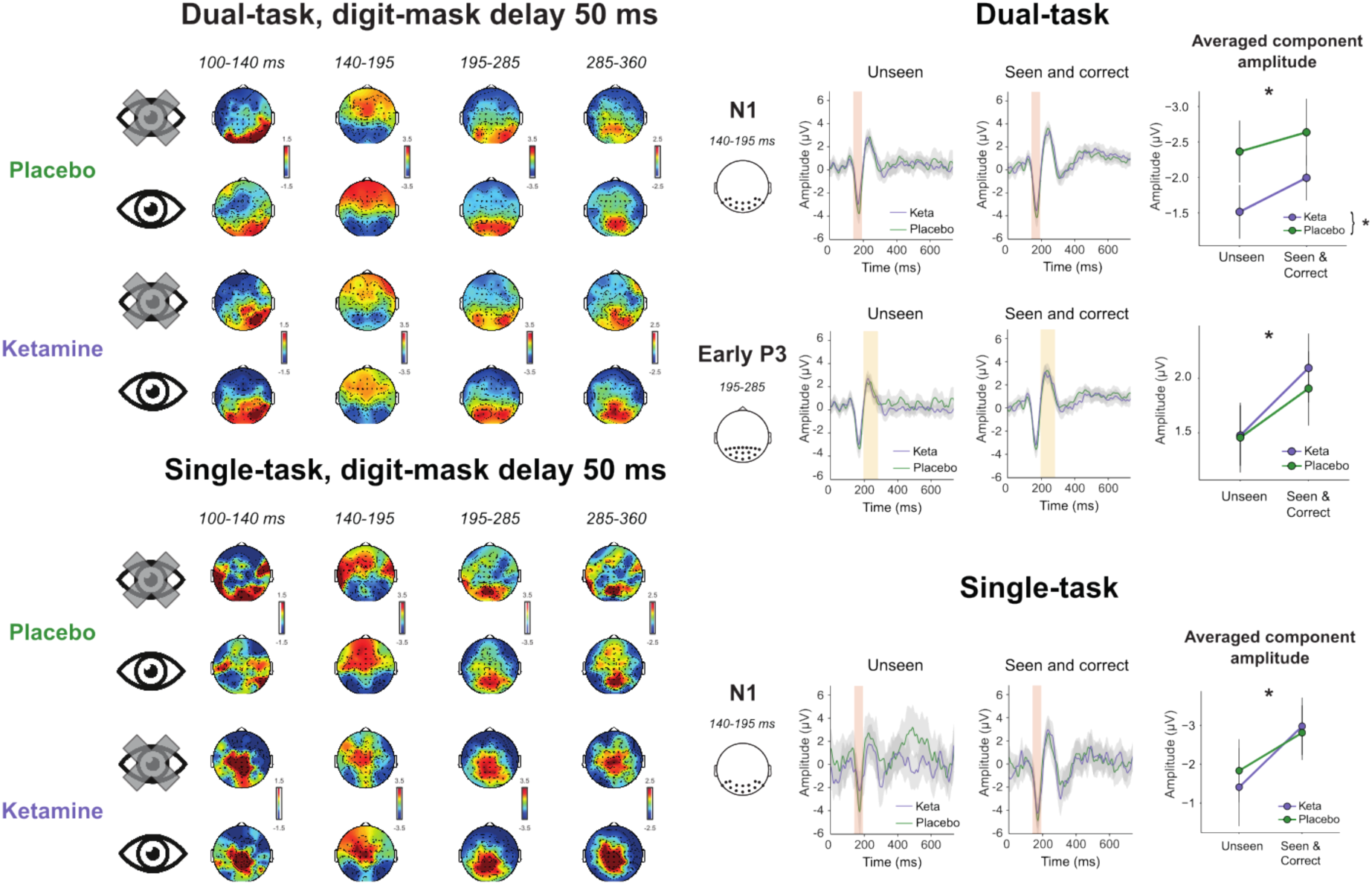
N1 tracks conscious access in both task contexts. Left. Mask-subtracted ERP topographies at the intermediate digit-mask delay (i.e., 50 ms) for each potential time-window (horizontally) in each pharmacological condition, i.e., placebo vs. ketamine, as a function of visibility, i.e., unconscious (crossed-out eye) vs. seen (vertically). Right. Subplots show the time-course of ERPs for the two pharmacological conditions (green: placebo, purple: ketamine). For each component, the preselected cluster of electrodes is depicted by black dots in the topographies at left. Preselected time-windows of interest, used for statistical analysis, are shown by colored rectangles. Grey shaded area around the curves represents one standard error of the mean. The line-plots on the right depict the averaged amplitude of each component in this window, i.e., across the corresponding cluster of electrodes, as a function of conscious access (i.e., seen and correct vs. unseen) and pharmacological condition. Error bars represent one standard error of the mean. * means p < 0.05 (top of the plot: seen and correct vs. unseen, legend: ketamine vs. placebo).

In the dual-task condition, conscious access positively correlated with N1 (*F_1,17_* = 5.36, *p* = 0.033) and early P3 amplitudes (*F_1,17_* = 8.60, *p* = 0.009, see Figure 7, top). The N1 amplitudes were significantly decreased under ketamine across conscious and unconscious trials (*F_1,17_* = 4.51, *p* = 0.049), with no significant interaction between conscious access and pharmacological condition (*F_1,17_* = 0.17, *p* = 0.69). Ketamine did not modulate early P3 (*F_1,17_* = 0.096, *p* = 0.76).

**Figure 7.**
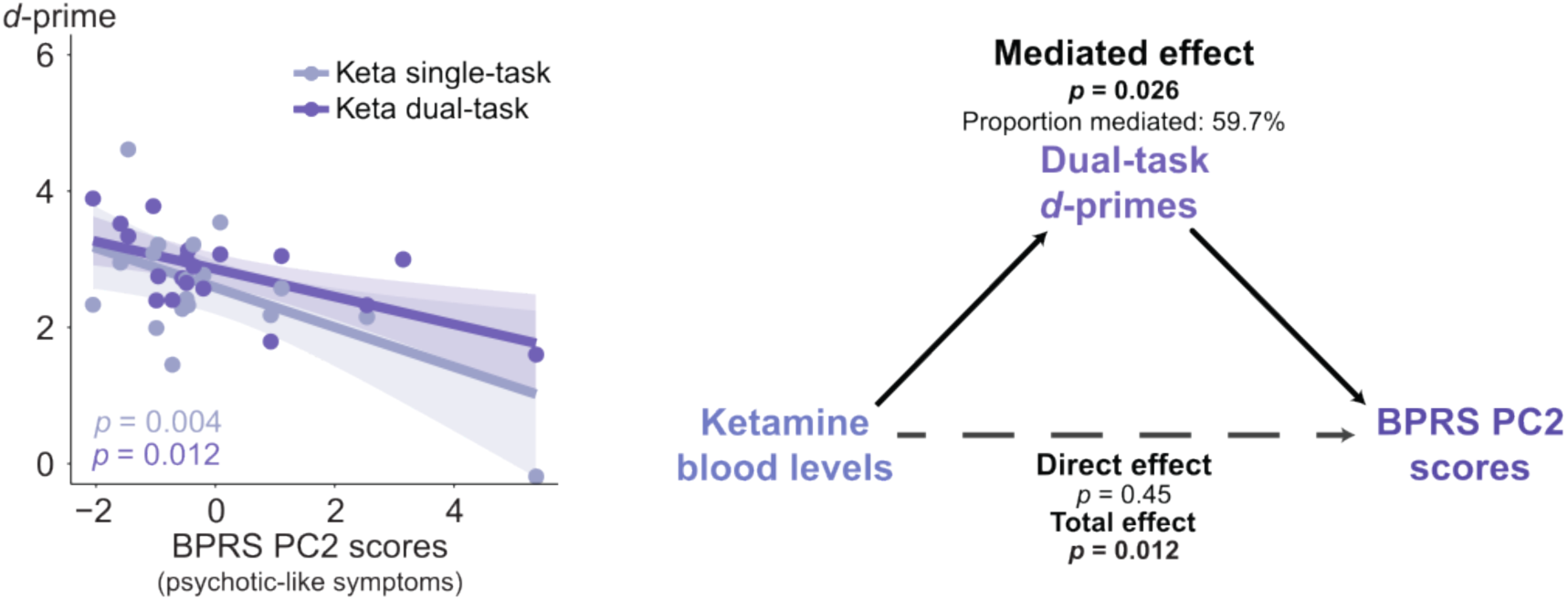
Relationships between consciousness disruption and psychotic-like symptoms. Left. Both in the single and the dual tasks, *d*-primes significantly correlated with BPRS PC2 scores. Each dot represents a participant, lines represent linear regressions, and shaded areas represent 95% confidence intervals. Right. Mediation analysis between ketamine blood levels, BPRS PC2 scores and dual-task *d*-primes. Ketamine blood levels significantly influenced the presence of BPRS PC2 scores capturing ketamine-induced psychotic-like symptoms. Effects mediated by dual-task *d*-primes were significant, while direct effects were not, suggesting that the influence of ketamine blood levels on psychotic-like symptoms intensity was fully mediated by detection impairments.

In the single-task condition, conscious access also positively correlated with the N1 amplitude (*F_1,16_* = 4.79, *p* = 0.044) without interaction with the pharmacological condition (*F_1,16_* = 0.41, *p* = 0.53, see Figure 7, bottom). No other significant effect was observed (all *p* > 0.1).

To sum up, N1 tracked conscious access in our task. The observation that ketamine intake is associated with an N1 decrease only in the dual-task condition suggests a compensation by attention on the effect of ketamine on sensory processing in the single-task condition. Moreover, the absence of interaction between pharmacological condition and visibility suggests that ketamine similarly decreases N1 in subliminal and conscious conditions.

### Relationships between consciousness impairments and ketamine-induced symptoms

#### Detection impairments are associated with psychotic-like symptoms

We finally investigated how ketamine-induced symptoms correlated with consciousness measures for each task. We found that impairments in detection *d*-primes were consistently associated with BPRS PC2 scores (dual-task condition: *F_1,17_* = 10.97, *p* = 0.004; single-task condition: *F_1,17_* = 7.82, *p* = 0.012; see Figure 7, left) but not with BPRS PC1 scores (all *p* > 0.1). There was no significant interaction between clinical scores and digit-mask delays either for detection or discrimination or significant effect of clinical scores on reaction times, performances on the sound, or ERP. Additional results can be found in the Supplementary materials.

#### Mediation analysis

To further explore the relationship between detection impairments and psychotic-like symptoms (i.e., BPRS PC2 scores), we conducted a mediation analysis. Given the correlation between ketamine blood levels and BPRS PC2 scores already described, we hypothesized that this relationship could be mediated by consciousness impairments.

We first ran a linear model confirming that ketamine blood levels significantly influenced the presence of psychotic-like symptoms (*t* = 2.23, *p* = 0.040, see Figure 1, B) and found that this effect was no more significant when dual-task detection *d*-primes were added in the model as a covariable (ketamine effect: *t* = 0.77, *p* = 0.45), while the main effect of *d*-primes was significant (*d*-prime effect: *t* = −2.26, *p* = 0.038). Therefore, we conducted a mediation analysis, which revealed that the correlation between ketamine blood levels and psychotic-like symptoms was mediated by decreased detection *d*-primes (ACME: CI = [0.001 0.03], *p* = 0.026; proportion mediated: 59.7%, CI = [−0.01 2.64], *p* = 0.055), with no direct effect (ADE: CI = [−0.011 0.03], *p* = 0.45, see Figure 7, right). A confirmatory analysis using a different approach to mediation analysis can be found in the Supplementary Materials.

Performing the same analyses with single-task *d*-primes was not conclusive because adding ketamine blood levels as a covariable rendered *d*-primes effects on psychotic symptoms below statistical significance threshold (*t* = −1.95, *p* = 0.69) precluding further exploration.

Overall, these results suggest that, in the dual-task condition, detection impairments mediate ketamine-induced psychotic-like symptoms.

## Discussion

### Summary of the results

In this study, we evaluated ketamine impact on conscious access in healthy volunteers using a randomized placebo-controlled cross-over within-subject paradigm and a cognitive task wherein conscious access was modulated both by masking and attentional distraction, while brain activity was measured using high-density EEG. At the behavioral level, we observed an increased effect of masking both for discrimination and detection. The psychological refractory period, reflecting dual-task interference, was also longer under ketamine compared to placebo. Ketamine disrupted conscious access through a reduction of early and late EEG components. Specifically, brain activity was significantly modulated by digit-mask delays; and ketamine altered N1 in the dual-task and the unattended condition, as well as early P3 in the single-task condition. Importantly, we found that N1 tracked conscious access as its amplitude was reduced in unconscious (i.e., unseen) compared to conscious trials (i.e., seen and correct). Finally, participants reported ketamine-induced psychotic-like symptoms which positively correlated with visual detection deficits. A mediation analysis suggested that dual-task detection alterations mediated the influence of ketamine blood levels on psychotic-like symptom intensity.

### Mechanisms of consciousness disruption by ketamine

Overall, our behavioral results suggest that low doses of ketamine disrupt conscious access through alterations of both sensory and attentional processing. This was particularly the case at the intermediate digit-mask delay, suggesting an elevation of consciousness threshold under ketamine. By contrast, there was no performance impairments at the longest digit-mask delay, indicating that conscious processing of clearly visible stimuli is largely preserved under ketamine.

In our paradigm, consciousness was tracked by N1 amplitude both in the single and the dual tasks. This result adds to the debate regarding the neural correlates of consciousness. While the P3 component was found to be a signature of conscious access in many studies^30,40–45^, empirical data suggest that it may reflect post-perceptual processes^31,46–53^. In line with previous studies^31,48,50–52^, we observe that N1 increased with digit-mask delays both in the attended and the unattended conditions, whereas P3 was modulated by task relevance. Such a result aligns with the proposal that early components reflect an accumulation of sensory evidence that may precede and contribute to conscious access^2^.

Interestingly, the N1 decrease under ketamine was particularly observable in the dual-task condition. This result could reflect an impairment of bottom-up sensory processing that is partly compensated by attentional top-down amplification in the single-task condition. This is also supported by the correlation between N1 amplitude and conscious access in both the single and dual-task condition, but an effect of ketamine on N1 only in the dual-task condition. Surprisingly, N1 amplitude was decreased under ketamine compared to placebo even on seen and correct trials. This may indirectly reflect an additional metacognitive effect of ketamine where ambiguous at-threshold stimuli are more likely to be labeled as “seen”. Additionally, ketamine increased the psychological refractory period, suggesting that attentional processes and dual-task monitoring are also impeded. Overall, ketamine may cause bottom-up sensory processing impairments which could be exacerbated by a higher cognitive load (dual-task) impacting attentional availability.

Other studies involving ketamine intake in healthy volunteers evidenced deficits in learning and decision-making for attended conscious information without obvious sensory processing impairments^14,15,27,54^. Specifically, ketamine decreases the ability to monitor confidence^15^, to integrate conflicting information^14^ and to use prior knowledge to stabilize perception^27^.

Interestingly, consciousness has been conceptualized as a perceptual decision based on an unconscious sensory evidence accumulation, the slope of which would depend on the physical properties and the relevance of incoming stimuli^2,55,56^. The brain would combine accumulated sensory evidence with priors, and sample the obtained distribution to provide an unequivocal conscious percept. In this perspective, the elevation of the consciousness threshold induced by ketamine could derive from alterations in such computations and decision-making processes by NMDA receptor antagonism.

### Ketamine does not reproduce the consciousness disruption observed in schizophrenia

Ketamine is usually used as a pharmacological model of schizophrenia. Previous ERP studies found that ketamine reduced N2, P2, and P3 amplitudes, processing negativity and mismatch negativity, in particular during oddball paradigms^29^, and specifically decreased N170 during facial emotion processing^57^. These results highlight that it can induce broader alterations depending on the task, some of them being similar to that observed in schizophrenia.

In our study, consciousness impairments elicited by ketamine were quite distinct from that observed in persons with schizophrenia. In them, consciousness deficits are characterized by a dissociation between preserved subliminal processing and abnormal conscious processing, along with an elevation of consciousness threshold^1–3,58^. EEG recordings show various deficits, including anomalies in early ERPs, such as the auditory P50 or the visual P1^59,60^. Importantly, a similar visual masking paradigm comparing healthy controls and patients with schizophrenia found that the only component which was decreased in schizophrenia was the P3^2^. Moreover, in this study, no significant difference between patients and healthy controls was evidenced under unattended conditions^2^.

By contrast, in the current study, conscious processing under ketamine seems to be relatively preserved as no behavioral difference was robustly observed at the longest digit-mask delay. Additionally, we observed significant differences under unattended conditions contrary to what was observed in schizophrenia^2^. Finally, although P3 was altered during the single task as in schizophrenia, N1 was also consistently altered across all conditions.

In other words, while schizophrenia consciousness deficits seem to be primarily underpinned by top-down attentional and post-perceptual processing impairments, consciousness disruption induced by low-doses of ketamine appears to be attributable to bottom-up sensory processing abnormalities which are partly compensated by attentional amplification.

### Detection impairment as a mechanism for psychotic-like symptoms

In our study, some but not all ketamine-induced subjective effects were dose-dependent. Indeed, psychotic-like symptoms increased with ketamine blood levels whereas it was not the case for manic-like and dissociative symptoms. In both the single-task and the dual-task conditions, only psychotic-like symptoms positively correlated with visual detection impairments. Interestingly, this correlation equally occurred at all digit-mask delays.

To further explore the relationship between psychotic-like symptoms, detection impairments and ketamine blood levels, we ran a mediation analysis which revealed that detection impairments mediated the relationship between ketamine blood levels and psychotic-like symptoms. This result suggests that detection deficits may play a role in psychotic-like symptoms induced by ketamine. It confirms predictions and theoretical proposals made in previous work^1^, as well as empirical findings showing that elevation of consciousness threshold is associated with positive symptoms intensity in schizophrenia^4^ and mediates the link between long-distance brain anatomic dysconnectivity and the advent of psychotic symptoms both in schizophrenia and bipolar disorder^3^. Put differently, this finding supports the hypothesis according to which a disruption of consciousness increases the liability to delusions, regardless of the underlying neural mechanisms.

However, disruptions of consciousness have also been observed in neuropsychiatric disorders devoid of psychotic symptoms^4,61–63^. Therefore, such disruptions could be a contributing factor, albeit not a sufficient condition, to prompt delusions. Additional deficits notably of metacognitive nature may be crucial for perceptual abnormalities to translate into delusional ideas^4^. Indeed, if an elevated consciousness threshold may significantly decrease the amount of information entering consciousness, promoting fictive interpretations, the inability to reject implausible explanations may be necessary for these ideas to settle and spread^64,65^. Consistently, ketamine-induced decision-making deficits observed in previous studies may not only play a role in elevating consciousness threshold as proposed above, but also in turning abnormal perception into delusional beliefs^14,15,54^.

### Conclusion

In this study, we showed that low subanesthetic doses of ketamine induced consciousness disruption through both bottom-up sensory processing impairments and attentional interferences, and a N1 ERP component reduction. Interestingly, these alterations are not fully superimposable on consciousness deficits observed in persons with schizophrenia, although they could directly favor the advent of psychotic-like symptoms. Further studies are needed to confirm and extend these results, in particular for confirming the link between detection disruption and psychotic-like symptoms and for exploring how decision-making alterations induced by ketamine may relate to consciousness impairments.

## Methods

### Participants

A total of 21 adult participants (16 females, 5 males, aged 19-33 years old, mean = 25.24, SD = 4.2) was tested for both visits. Inclusion criteria are detailed in the Supplementary Materials. Three additional participants were included but were sick during the ketamine visit and could not perform all tasks. Other participants had to be excluded from behavioral (n = 2) and EEG (n = 1) analyses (see Supplementary Materials for details). Participants provided written informed consent and received a financial compensation for their participation in the study. The study received ethical approval from relevant authorities: the Agence Nationale de Sécurité du Médicament et des Produits de Santé and the Comité de Protection des Personnes Ile-de-France III (ID RCB: 2013-002056-33) and was registered under reference NCT02235012.

### Design and Procedure

#### Pharmacological protocol

We used a double-blind, placebo-controlled, randomized crossover design. Each participant came twice to the lab at least one month apart and received once ketamine (maximum 1 mg/kg over 2h) and once placebo (sodium chloride) in a randomized order to which both participants and experimenters were blind. Ketamine (or saline) was administered intravenously using a three-stage infusion protocol: a 10 min bolus stage (0.023 mg/kg/min), followed by a 20 min stabilization stage (0.0096 mg/kg/min) and a third open-ended stage (0.0048 mg/ kg/min) until task completion (∼60-90 min). The protocol was similar to previously published research^14^.

#### Visual perceptual task

The experimental paradigm is summarized in Figure 2. We used a variant of the masking paradigm designed for previous studies in healthy and clinical populations^2,3,36,37,41,62,63^. Our goal was to disrupt conscious access both through visual masking and attentional distraction.

Stimuli presentation began with a central fixation cross. A sound was played after a jittered delay (between 1000 and 1667 ms) to be unpredictable. The sound was an isolated syllable “*ka*” or “*pi*” pronounced by a female voice during 195 ms. After a first delay (sound-digit delay with a stimulus onset asynchrony, SOA, of 100, 300, or 500 ms, randomly intermixed across trials), a digit (2, 3, 7 or 8) was presented for a fixed duration of 17 ms above or below the fixation cross (randomly intermixed across trials). After a second delay (digit-mask delay with a SOA of 33, 50 or 183 ms randomly intermixed across trials), a metacontrast mask composed of four letters appeared at the digit location for 200 ms. Twenty percent of the trials were mask-only trials (catch trials): the digit was replaced by a blank screen of the same duration, i.e., 17 ms.

The exact same sequences of stimuli were presented under three distinct conditions, which differed only in the instructed task. The dual-task instructions were to pay attention both to the sound and to the masked digit, and to give three behavioral responses in the following order: (1) determining as fast as possible whether the sound was “*ka*” or “*pi*”, (2) deciding as fast as possible whether the digit was greater or smaller than 5 (which provided an objective measure of the digit perception) and (3) reporting the digit visibility using a categorical response “Seen” or “Unseen” (providing a subjective measure of conscious access). When performing the single task, participants had to answer questions about the digit only, i.e. (2) and (3), and were asked to ignore the sound. Finally, in the unattended condition, participants had to answer the question about the sound only, i.e. (1), and ignoring the digit.

Participants provided answers to the digit and the sound objective questions by pressing as fast as possible specific keys of a keyboard. The keys for the visibility question were randomized on a trial-by-trial basis (see details in the Supplementary Materials).

Subjects completed eight blocks in total: six “dual-task” blocks of 90 trials each (i.e., 540 trials) that were performed in a row, one “single-task” block of 90 trials, and one “unattended” block of 90 trials. In the dual-task condition, there were 60 trials (including 12 mask-only trials) in each combination of sound-digit delay (3 levels, i.e., 100, 300, or 500 ms) and digit-mask delay (3 levels, i.e., 33, 50, 183 ms). Under the single-task and the unattended conditions there were 10 trials, including 2 mask-only trials, in each combination of sound-digit and digit-mask delays.

Our experiment was coded with *MATLAB* using *Psychtoolbox* functions^66,67^.

#### Clinical measures and ketamine dosages

Participants underwent psychiatric symptom measurements at each visit 30 minutes after the infusion started right before the tasks began. The measurements consisted of the 24-item Brief Psychiatric Rating Scale (BPRS)^68^ assessing general psychiatric dimensions, and the 23-item Clinician-Administered Dissociative States Scale (CADSS)^69^ measuring “dissociative” symptoms. Participants also underwent physical measurements (blood pressure, heart rate, blood oxygen saturation) before infusion and every 15–20 min. The level of ketamine plasma concentration was controlled by a blood sample taken 30 minutes after infusion start, which was immediately centrifuged for 10 min at 1700 × g and 4 °C using a Multifuge 3 S centrifuge (Fisher Scientific SAS). The resulting plasma was frozen at −20 °C and sent to the pharmacology lab to be analyzed. Ketamine plasma concentration was later estimated using a Waters 600 high-performance liquid chromatography system and a Waters 2996 photodiode array detector (Waters Corporation).

#### EEG

##### Recording

EEG data were recorded using a 64-channel BioSemi ActiveTwo system (BioSemi, Amsterdam, The Netherlands) at a sampling frequency of 1024 Hz. The 64 scalp electrodes were positioned according to the 10/20 system.

##### Preprocessing

The raw EEG data were down-sampled to 216 Hz, low-pass filtered at 40 Hz, high-pass filtered at 0.1 Hz and notch filtered at 50, 100 and 150 Hz. EEG data were epoched at the trial level. Individual epochs were visually inspected to remove epochs containing large jumps, and to identify bad electrodes whose activities were reconstructed using spherical interpolation based on neighboring electrodes. Remaining stereotyped artifacts (e.g., eye blinks) were removed using an Independent Component Analysis (ICA) and identifying eye blink components manually for each participant and each condition (i.e., visits and tasks). Following the removal of ocular components, data were re-epoched to selectively explore the digit processing (digit-locked, −1 to 1.4 s relative to the digit onset) and a second visual inspection was performed to eliminate bad trials or electrodes. Digit-locked trials were corrected for baseline over a 400 ms window during fixation at the beginning of each trial. These preprocessing steps were performed using the *FieldTrip* toolboxes for *MATLAB* and additional custom-written scripts.

##### Event-related potentials

Digit-locked trials were separated into digit and catch trials that were subsequently averaged for each delay (sound-digit and/or digit-mask), task and visits at the subject level. The averaged activity of catch trials was subtracted from that of digit trials for each condition (delays, task, visit), to isolate the digit-evoked activity^2,41^. ERP latency highly varied according to the digit-mask delays, so different time-windows were used for each delay based on visual observation of EEG signals averaged across subjects and pharmacological conditions for each digit-mask delays in the dual-task condition (see Figure 4). Electrodes clusters were defined based on topographies for each time-windows. For each subject, under each condition of interest, the EEG signals were then averaged over clusters of electrodes and time-windows.

To further explore the neural correlates of consciousness, we run additional analyses on digit-locked ERP separating conscious (rated as “Seen” and correctly compared to 5) versus unconscious trials (“Unseen”) at the intermediate digit-mask delay (50 ms).

### Statistical analyses

All statistical analyses were two-tailed and performed using the *R* statistical software (https://www.r-project.org). They were not adjusted for multiple comparisons.

For behavioral measures, analyzes of variance (ANOVAs) and Pearson correlations were conducted after excluding extreme reaction times (shorter than 50 ms or longer than 2.5 seconds for the sound task, shorter than 200 ms or longer than 2.5 seconds for the digit discrimination task) with digit-mask delay and/or sound-digit delay, task and pharmacological condition as within-subject factors. For reaction times analyses in the dual task, only trials on which participants correctly answered the sound task were taken into account.

For the ERP, ANOVAs were conducted separately for each ERP components on their corresponding averaged amplitude (over electrodes and time points), with digit-mask delay, and/or sound-digit delay, task and pharmacological condition as within-subject factors.

Since BPRS is a heterogeneous scale, we run a principal component analysis (PCA) on all the BPRS items using the same approach and analytical tools as in previous studies^4,70–73^.

Principal components (PC) capture how the symptoms are correlated (or anticorrelated) one with another, thereby reflecting fully data-driven symptomatic dimensions, or clinical geometry. Each item was first scaled to have unit variance across participants before running the PCA (items having null variance are discarded). Significance of the derived PC was computed via permutation testing. For each permutation, participant order was randomly shuffled for each symptom variable before re-computing PCA. This permutation was repeated 1,000 times to establish the null model. PCs which accounted for a proportion of variance that exceeded chance (*p* < 0.05 across all 10,000 permutations) were retained for further analysis and ordered as a function of the proportion of symptom variance they account for.

Finally, we examined the link between detection impairments, psychotic-like symptoms (i.e., BPRS PC2 scores) and ketamine blood levels using a mediation analysis^74,75^. Two linear models (*d*-primes as a function of ketamine blood levels and psychotic-like symptoms as a function of ketamine blood levels and *d*-primes) in a mediation analysis with 10,000 simulations, using the mediate function included in the R software mediation package^76^.

## Acknowledgments

We thank all the participants and the team of the « Centre pour la Recherche Clinique » (CRC) of Sainte Anne hospital. LB thanks the Fondation Bettencourt-Schueller, the Philippe Foundation, the Foundation L’Oréal-Unesco and the National Institute of Mental Health (R01MH116038 and U01MH121766) for their support.

