## Supplementary Materials for "Ketamine disrupts consciousness in healthy participants in relation with psychotic-like symptoms"

### Supplementary methods

#### *Participants*

Inclusion criteria were the following: right-handed, normal or corrected-to-normal vision. Non-inclusion criteria included: history of neurological or psychiatric disease, family history of psychotic disorders, addiction to psychoactive drugs, history of psychotropic medication, history of cardiac disease, blood pressure above 140/90 mmHg, pregnancy or breastfeeding.

Two participants had to be excluded from the behavioral analyses because of they had abnormal results. One performed the wrong single-task under placebo (answered about the sound category instead of the digit and then rated the digit visibility) and had abnormally low visibility levels at the intermediate digit-mask delay (less than 25% of trials were seen). One participant had an atypical pattern of response with poor performances for discrimination, and a high discrepancy between discrimination and visibility ratings, suggesting that she may have misunderstood the instructions. One additional participant had to be excluded from the EEG analyses because the recording during one of the two visits was unexploitable. One of the remaining participants had no EEG “unseen” trials under placebo at the intermediate digit-mask delay (50 ms) in the single task and was therefore excluded from the analyses investigating the EEG neural correlates of conscious access for this task only.

#### *Design and Procedure*

*Pharmacological protocol*
For the ketamine visit, a preparation of racemic ketamine 0.1% was assembled using 2 mL of a 5 mL phial of injectable Panpharma ketamine 250 mg/5 mL (containing 288.4 mg of ketamine chlorhydrate, corresponding to 250 mg of base ketamine), which was added to a 100 mL bag of saline solution (Macropharma sodium chloride 0.9%), of which 2 mL had been extracted. For the placebo visit, the 100 mL bag of saline solution was used *ad integrum*. The ketamine and placebo bags were indistinguishable. The pump used was a programmable Volumat Agilia pump (Fresenius Kabi France SAS), programmed with the above protocol and which switched sequentially from one stage to the next automatically.

*Visual perceptual task*
Participants provided answers to the digit and the sound objective questions by pressing as fast as possible specific keys of a keyboard. For each participant, one hand was dedicated to the sound discrimination task, the other to the digit comparison task. The answer “*smaller than 5*” and “*ka*” were always assigned to the left most button for each hand. In the dual-task condition, participants were instructed to perform the sound discrimination task first and as

quickly as possible without waiting for the digit to appear. In the dual and the single tasks, as soon as participants had responded to the digit objective question (or after five seconds in the absence of response), a screen for the visibility question appeared. The response words "Vu" ("Seen") and "Non Vu" ("Unseen") were displayed on the screen, randomly assigned to the right and left of the fixation point. Participants responded by pressing whatever buttons on the side of the response they wanted to select (e.g., using left hand for "Vu" if it was presented on the left of the fixation cross). The mapping between the keys and the response options was randomized on a trial-by-trial basis to decouple participants' responses to the objective questions from the response to the visibility question. No time limit was assigned to the visibility question and the "Seen" and "Unseen" response choices remained on the screen until a response was given.

Participants were trained to perform the three tasks during the screening visit and at the beginning of each visit. To facilitate learning, the training block order was the same for all participants: first they performed the unattended block (i.e., sound discrimination task), then the single (digit-related) task and finally the dual task, so that complexity progressively increased. They did at least 20 trials of each task before starting the experiment, and training continued until performances reached a ceiling.

### Supplementary results

#### *Ketamine-induced symptoms*

There was a significant positive correlation between CADSS and BPRS total scores ($r = 0.51$, $t_{19} = 2.57$, $p = 0.019$). This was driven by a positive correlation between CADSS and the manic-like dimension of BPRS, i.e., PC1 ($r = 0.62$, $t_{19} = 3.47$, $p = 0.003$), but not with the psychotic-like dimension of BPRS, i.e. PC2 ($r = -0.41$, $t_{19} = -1.95$, $p = 0.067$).

#### *Behavioral measures*

*Attentional interferences*
RT increase as a function of digit-mask delay ($F_{2,36} = 5.53$, $p = 0.008$) with no interaction with the pharmacological condition or with the sound-digit delay (all $p > 0.3$).

The PRP effect size usually depends on the RT in performing the task on the first stimulus (the longer the first RT, the higher the slowdown for the second RT). When splitting the trials according to "slow" or "fast" RT to the sound (for each subject, under and above their own median), we indeed observed a main effect of this parameter on the digit RT ($F_{1,18} = 66.61$, $p < 0.001$). This parameter significantly interacts with the sound-digit delay ($F_{2,36} = 41.93$, $p < 0.001$) and with the pharmacological condition ($F_{1,18} = 8.41$, $p = 0.010$) but there was no triple interaction between these three variables ($F_{2,36} = 0.98$, $p = 0.39$).

There was no classical attentional blink effect. In the dual task, detection was not significantly modulated by sound-digit delays ($F_{2,36} = 3.12$, $p = 0.056$) but there was a significant interaction between sound-digit delay and the pharmacological condition ($F_{2,36} = 4.61$, $p = 0.017$). This effect corresponds to a decrease in discrimination as a function of sound-digit delays under placebo ($F_{2,36} = 6.45$, $p = 0.004$) that disappeared under ketamine ($F_{2,36} = 0.51$, $p$

= 0.60). Regarding detection *d*-primes no significant effect of sound-digit delays was observed.

*Poorer performances in the sound-related task*
Participants were overall less able to perform the sound-related task under ketamine. In terms of discrimination performance, there was a main effect of ketamine across the dual task and the single sound-related task driven by an impairment in the single sound-related task (ketamine effect: $F_{1,18} = 5.06$, $p = 0.037$; task effect: $F_{1,18} = 2.42$, $p = 0.14$; interaction task × ketamine: $F_{1,18} = 4.27$, $p = 0.053$; single task sound: ketamine effect: $t_{18} = 2.33$, $p = 0.032$; dual task: $t_{18} = 1.04$, $p = 0.32$).

Participants were also much slower in performing the sound task under ketamine in both the single sound-related task and the dual-task conditions (ketamine effect: $F_{1,18} = 19.17$, $p < 0.001$; task effect: $F_{1,18} = 46.31$, $p < 0.001$; interaction task × ketamine: $F_{1,18} = 4.75$, $p = 0.043$; single task sound: ketamine effect: $t_{18} = -4.02$, $p < 0.001$; dual task: $t_{18} = -3.8$, $p = 0.001$).

***EEG measures***

*Sound-digit delays effects on EEG components*
In the placebo condition, sound-digit delays had only few effects on EEG components. In the dual task, there was no main effect of this parameter on any component (all $p > 0.05$). However, the interaction between the sound-digit and the digit-mask delays was significant for N2 ($F_{2,34} = 5.26$, $p = 0.010$) and close-to-significance for early P3 ($F_{2,34} = 3.27$, $p = 0.050$). In the single task, no effect or interaction involving sound-digit delays were significant ($p > 0.1$). In the single sound-related task, the only significant result was a main effect of sound-digit delays for the late P3 ($F_{2,34} = 3.80$, $p = 0.032$) reflecting an increased amplitude at the intermediate sound-digit delay (300 ms).

When examining ketamine effects, the only interaction between the pharmacological condition and the sound-digit delays was found in the unattended condition for the late P3 ($F_{2,33} = 5.47$, $p = 0.0089$), which overall exhibited an opposite pattern in comparison to the placebo condition (i.e., was increased rather than increased for sound-digit delay of 300 ms). There was no triple interaction between the pharmacological condition, the sound-digit delays and the digit-mask delays (all $p > 0.1$).

***Relationships between consciousness impairments and ketamine-induced symptoms***

*Detection impairments are associated with psychotic-like symptoms*
In the dual-task condition, we found that detection *d*-primes significantly decreased as a function of BPRS PC2 scores, i.e., psychotic-like symptoms ($F_{1,17} = 10.97$, $p = 0.004$, see Figure 7, left) but nor BPRS PC1 scores, i.e., manic-like symptoms ($F_{1,17} = 0.12$, $p = 0.73$, 1/BF = 3.63) neither BPRS total scores ($F_{1,17} = 0.83$, $p = 0.38$, 1/BF = 3.13). Surprisingly, CADSS scores also varied with detection *d*-primes but the higher the dissociative symptoms, the better the performance ($F_{1,17} = 9.78$, $p = 0.006$). By contrast, discrimination performances were not modulated by any clinical scales. There was no significant interaction between clinical scores and digit-mask delays either for detection or discrimination (all $p > 0.1$).

In the single-task condition, BPRS PC2 scores were also associated with decreased detection *d*-primes ($F_{1,17} = 7.82$, $p = 0.012$, see Figure 7, left). There was no significant effect of other clinical measures (BPRS total scores: $F_{1,17} = 4.09$, $p = 0.059$; all other $p > 0.1$) or their interaction with digit-mask delays (all $p > 0.1$). There was no significant effect of clinical scores on discrimination (BPRS PC2 scores: $F_{1,17} = 4.32$, $p = 0.053$; all other $p > 0.1$).

No significant effect of clinical scores on reaction times or on performances on the sound was observed (all $p > 0.05$).

No interaction between EEG measures and clinical scores was found.

*Mediation analysis*

As a confirmatory analysis, we correlated the residuals of two linear models: 1) *d*-primes as a function of ketamine blood levels, 2) BPRS PC2 scores as a function of ketamine blood levels. We found that residuals were significantly correlated ($t_{17} = -2.33$, $p = 0.032$), supporting that a statistical relationship exists between *d*-primes and BPRS PC2 scores when removing ketamine blood levels influence.
